# Cardiovascular responses to antigravity muscle loading during head-down tilt at rest and after dynamic exercises

**DOI:** 10.1101/399477

**Authors:** Cristiano Alessandro, Amirehsan Sarabadani Tafreshi, Robert Riener

## Abstract

The physiological processes underlying hemodynamic homeostasis can be modulated by muscle activity and gravitational loading. The effects of antigravity muscle activity on cardiovascular regulation has been observed during orthostatic stress. Here, we evaluated such effects during head-down tilt (HDT). In this posture, the gravitational gradient along the body is different than in upright position, leading to increased central blood volume and reduced venous pooling. We compared the cardiovascular signals obtained with and without antigravity muscle loading during HDT in healthy human subjects, both at rest and during recovery from leg-press exercises. Further, we compared such cardiovascular responses to those obtained during upright position. We found that loading the antigravity muscles during HDT at rest led to significantly higher values of arterial blood pressure than without muscle loading, and restored systolic values to those observed during upright posture. Maintaining muscle loading post-exercise altered the short-term cardiovascular responses, but not the values of the signals five minutes after the exercise. These results demonstrate that antigravity muscle activity modulates cardiovascular regulation during HDT. This modulation should therefore be considered when interpreting cardiovascular responses to conditions that affect both gravity loading and muscle activity, for example bed rest or microgravity.

## 1 Introduction

The autonomic nervous system can accomplish hemodynamic homeostasis under a variety of conditions, including during physical exercise and gravitational stress. During orthostatic challenges, for example, sympathetic vasoconstrictor activity maintains blood pressure, avoiding postural hypotension and syncope [51, 52]. While such activity is primarily regulated by baroreflexes, previous research demonstrated that contraction of the antigravity muscles modulates cardiovascular regulation [48, 29, 11, 18, 55]. Similarly, other works showed that antigravity muscle activity influences the physiological processes involved in recovery from physical training during orthostasis [22, 23]. Whether these findings hold true also in the absence of orthostatic stress is still unclear. A posture like head-down tilt (HDT, i.e. subjects lie supine on a head-down tilted platform), for example, increases central blood volume (CBV), stimulates baroreceptors and facilitates venous return (Fig. 1a-b). These factors modulate the mechanisms of hemodynamic homeostasis [56, 24, 32, 42], and so may also modify the role of antigravity muscle activity on cardiovascular function.

**Figure 1:**
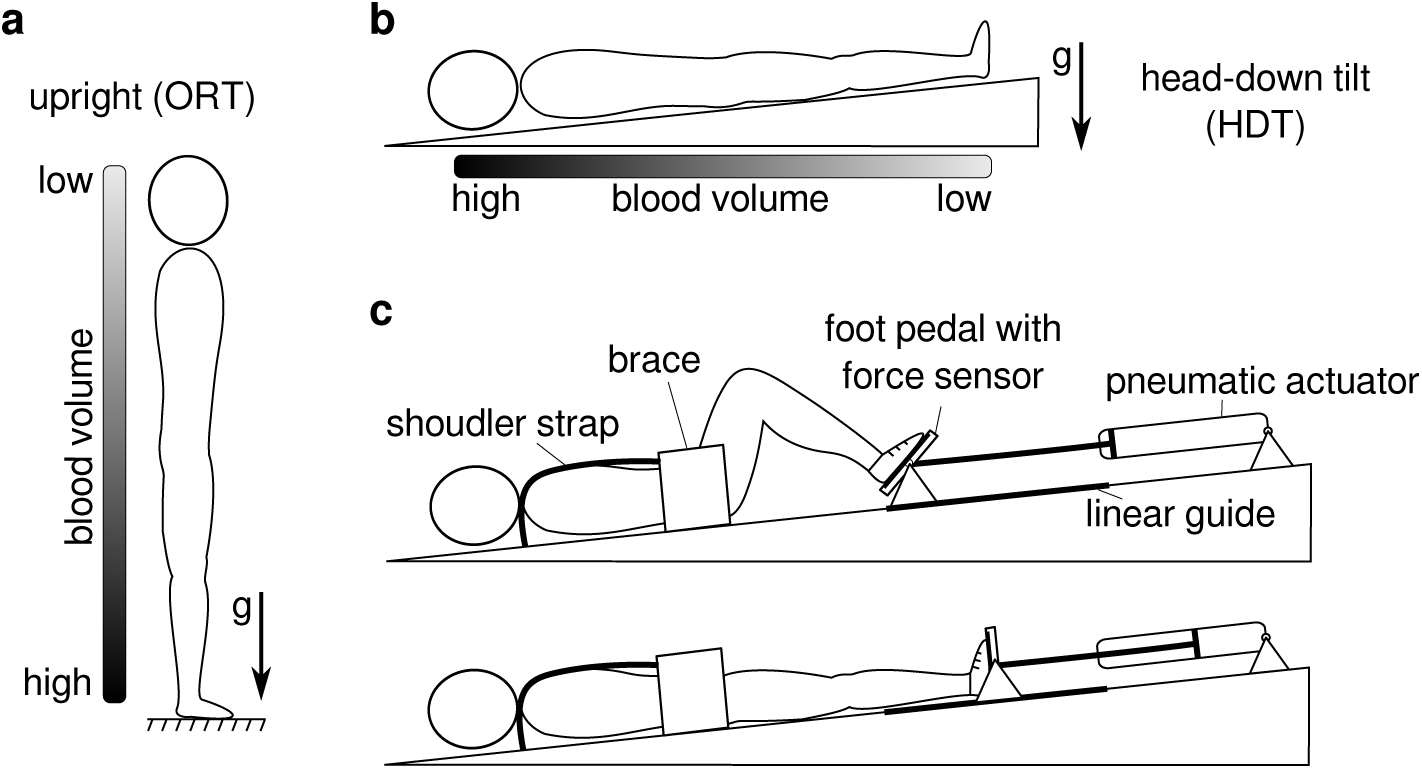
Upright and head-down tilt postures, and schematic representation of the robotic leg-press device. During upright posture (A), blood tends to shift towards the lower part of the body under the effect of gravity. This displacement of blood is regulated by physiological mechanisms that accomplish hemodynamic homeostasis, which are modulated by the activity of the antigravity muscles. During head-down tilt (B), gravity leads to a displacement of blood towards the upper part of the body, and since there is no weight bearing, antigravity muscles are inactive. The robotic leg-press device MARCOS (C) allowed us to apply antigravity muscle loading and perform leg-press exercises during HDT. The device provides simulated ground reaction force by means of pneumatic actuators that oppose leg extension movements, and maintains the desired force using force feedback loops that receive force measurements from the foot pedals. Additional details on this device are described in Sec. 4.3.

Head-down tilt is extensively used in the context of space physiology to simulate the effects of microgravity on the cardiovascular system [25, 37, 34, 17]. Experiments have shown that short-term exposure to HDT reduces heart-rate, mean arterial pressure, and leg blood volume, and increases cardiac output [50, 31, 37, 30]. In the long term, HDT induces a decrease of plasma volume, reduces baroreflex sensitivity, and may ultimately cause orthostatic intolerance [34]. Compared to upright posture, however, HDT (and in general any supine position) not only removes orthostatic stress, but also unloads the antigravity muscles (Fig. 1a-b). The observed cardiovascular responses may therefore be due to a combination of these two factors. To better interpret these cardiovascular responses, it is therefore important to determine the potential role of antigravity muscle activity on cardiovascular function. Establishing this role may also contribute to a better understanding of the cardiovascular deconditioning caused by microgravity [20, 16, 53, 7, 57, 8], and of the physiological processes involved in recovery from physical exercises during space missions [15, 54].

In this study, we evaluated the effects of antigravity muscle loading during HDT. To do so, we developed a head-down tilted platform equipped with a robotic leg-press device [19], which allowed us for the first time to apply antigravity muscle loading during HDT (Fig. 1c). We collected systolic, diastolic, mean blood pressure, pulse blood pressure and heart rate from healthy volunteers lying on the tilted platform, at rest as well as immediately after they performed bouts of leg-press exercises. We performed these recordings both with muscle loading (HDT-ML), when subjects maintained their legs extended against the resistance of the device, and without muscle loading (HDT-noML), when subjects maintained their legs extended against no external resistance (i.e. the device did not apply any force). Additionally, we recorded the cardiovascular signals during orthostatic stress (i.e. upright posture at rest), allowing us to compare these signals to the cardiovascular responses observed during HDT.

We found that muscle loading during HDT modified mainly the cardiovascular signals at rest, and only minimally affected their short-term dynamics after exercises. The blood pressure values obtained with muscle loading at rest were significantly higher than those obtained without muscle loading, and were close to those observed during orthostatic stress. This observation suggests that the cardiovascular responses to HDT are partially due to the lack of antigravity muscle activity, and not only to the different gravitational gradient. These results are consistent with the hypothesis that antigravity muscle activity modulates cardiovascular functions in the absence of orthostatic stress.

## 2 Results

### 2.1 Head-down tilt without antigravity muscle loading

We first evaluated the cardiovascular responses to head-down tilt without antigravity muscle loading (HDT-noML) at rest (i.e. subjects laid on the tilted platform without performing any movement). Figure 2 illustrates an example of the cardiovascular signals for a representative subject. HDT-noML caused a reduction of the entire blood pressure (BP) trace (i.e. lower values of both systolic and diastolic BP), a minimal increase of pulse BP (pBP), and a very evident decrease of heart rate (HR).

**Figure 2:**
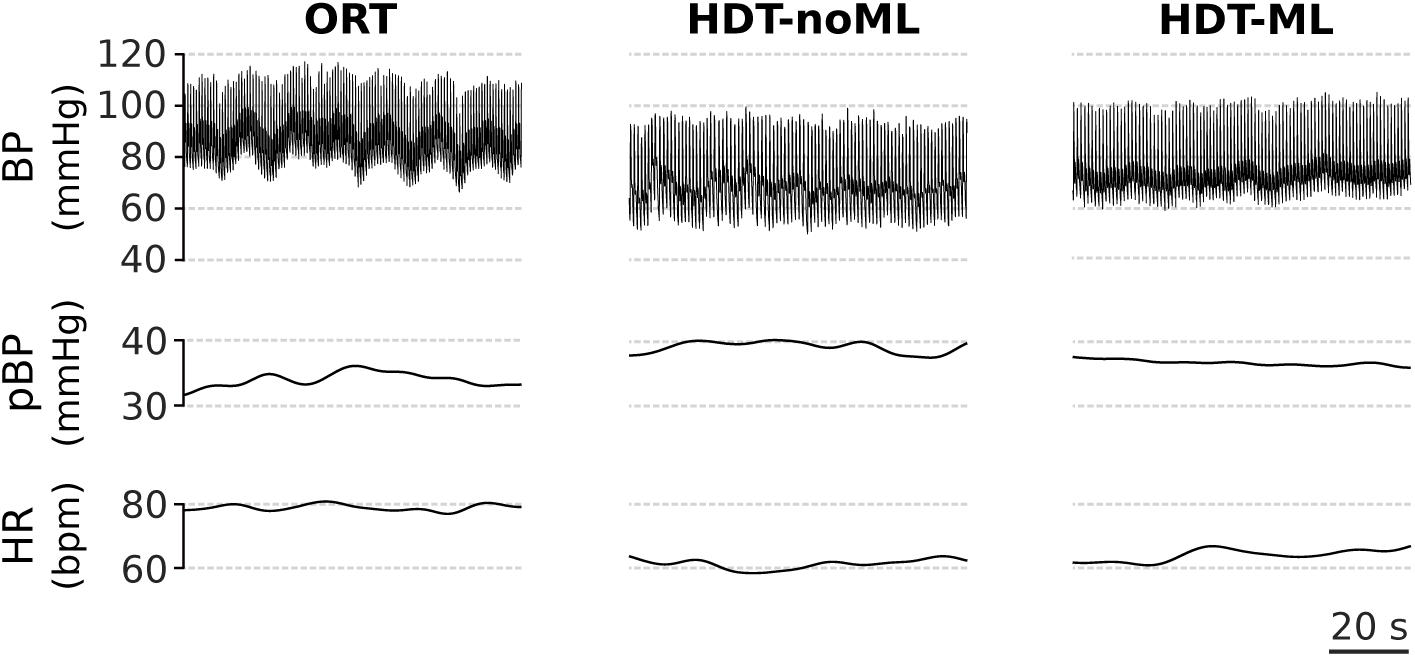
Cardiovascular signals at rest for a representative subject. During head-down tilt without antigravity muscle loading (HDT-noML), the overall BP trace (i.e. systolic and diastolic) was lower than during both orthostatic stress (ORT) and head-down tilt with muscle loading (HDT-ML). In the HDT posture, pulse BP was slightly higher and heart rate was lower than during ORT, and they were minimally affected by antigravity muscle loading.

These results were consistent across subjects, as shown in Fig. 3. During HDT-noML, systolic (sBP), diastolic (dBP) and mean blood pressure (MAP) were significantly lower than during orthostatic stress (ORT; sBP: –14.9±2.9 mmHg, p<0.001; dBP: –17.8±2.6 mmHg, p<0.001; MAP: –16.9±2.5 mmHg, p<0.001); pulse BP was slightly higher but not significantly different (+3.1±2.2 mmHg, p=0.321); and HR was significantly lower (–19.8±2.1 bpm, p<0.001).

**Figure 3:**
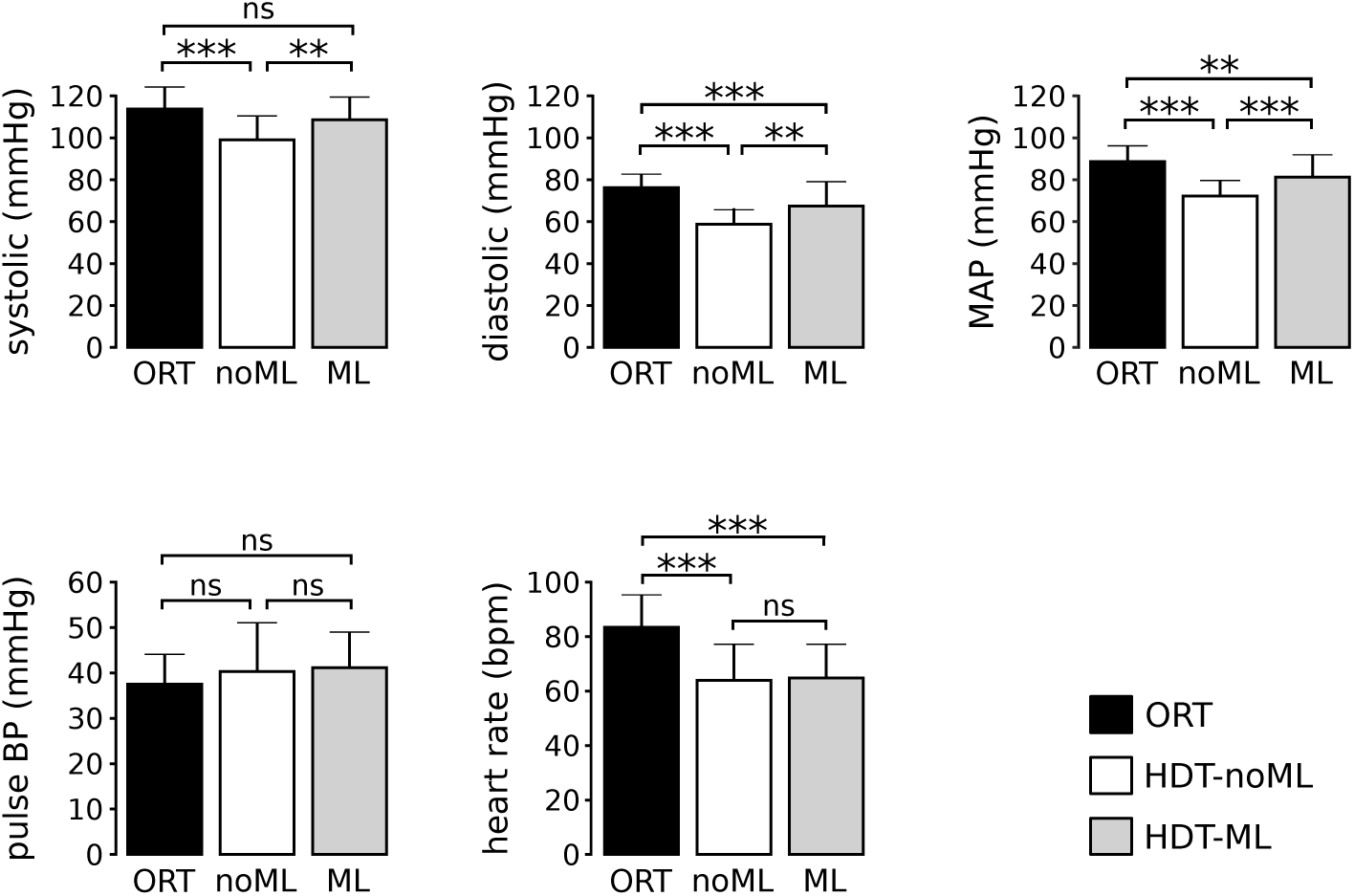
Cardiovascular signals at rest for all subjects. During head-down tilt without muscle loading (HDT-noML), systolic BP, diastolic BP, mean arterial pressure and heart-rate were lower than during orthostatic stress (ORT). During head-down tilt with muscle loading (HDT-ML), systolic BP, diastolic BP and mean arterial pressure were higher than during HDT-noML, while pulse BP and heart rate were not different from HDT-noML. Note that systolic BP during HDT-ML was not different from ORT. Data are presented as mean±s.d. across subjects. N=16, 17, 16 subjects contributed to the bars ORT, HDT-noML, and HDT-ML respectively for all signals (see Sec. 4). ***p<0.001, **p<0.01, *p<0.05.

### 2.2 Antigravity muscle loading during HDT at rest

Compared to orthostatic stress (ORT), HDT without muscle loading (HDT-noML) both altered the gravitational gradient along the body and removed weight bearing, hence unloading the antigravity muscles. To determine the relative contributions of these two changes on the observed physiological measures, we applied antigravity muscle loading during HDT (condition HDT-ML) by means of a robotic leg-press device (see Sec. 4.3), and compared the obtained cardiovascular responses to those observed during ORT and HDT-noML. Such analyses were instrumental to: (1) evaluate the influence of antigravity muscle activity on cardiovascular functions during HDT, and (2) assess the extent to which the results obtained during HDT-noML (Sec. 2.1) were driven by the lack of antigravity muscle activity.

We found that antigravity muscle loading increased arterial blood pressure during HDT at rest, and restored similar values to those observed during orthostasis, as illustrated in Fig. 2 and 3. The HDT-ML condition induced significantly higher values of systolic BP (+9.5±2.8 mmHg, p=0.002), diastolic BP (+8.7±2.5 mmHg, p=0.002) and mean arterial pressure (+8.9±2.4 mmHg, p<0.001) compared with those elicited by HDT-noML. In contrast, it resulted in non-significantly different values of pulse PB (+0.7±2.2 mmHg, p=1) and heart rate (+1.4±1.3 mmHg, p=0.92) compared to HDT-noML. Such higher values of systolic BP were not significantly different from ORT (–5.4±2.9 mmHg, p=0.132), unlike what was observed without loading the antigravity muscles. On the other hand, the HDT-ML values of diastolic BP, MAP and heart rate were significantly lower (dBP: – 9.1±2.6 mmHg, p<0.001; MAP: – 7.8±2.5 mmHg, p=0.003; HR: – 18.6±2.1 bpm, p<0.001), and pulse BP was not significantly different from ORT (+3.8±2.2 mmHg, p=0.175), consistent with what was observed during HDT-noML. These results indicate that antigravity muscle activity modulates the mechanisms of blood pressure regulation during HDT at rest, and suggest that such muscle activity contributed to the BP differences between ORT and HDT-noML described above (Sec. 2.1).

### 2.3 Antigravity muscle loading during HDT after leg exercise

Antigravity muscle activity may also affect the cardiovascular processes involved in recovery from dynamic exercises during HDT. We investigated this issue by comparing the cardiovascular signals obtained after performing dynamic leg-press exercises in two scenarios: (1) when antigravity muscle loading was applied before and after the exercise bouts, and (2) when muscle loading was not applied.

Figure 4 illustrates an example of the the cardiovascular signals obtained without (HDT-noML) and with (HDT-ML) antigravity muscle loading for one subject. During the first 90s of recovery (early recovery, R_*early*_), the signals exhibit clear transient dynamics, which disappear within 200s when cardiovascular processes have stabilized. During early recovery, diastolic BP and MAP feature well evident undershoots with respect to baseline (i.e. value of the signals before the exercise bout, BAS), and then they stabilize at values slightly lower than baseline for this subject. Systolic BP follows a similar trend, but with a less clear minimum peak. Pulse BP keeps increasing at the beginning of the recovery phase, and then it returns to values similar to baseline. Finally, heart rate slowly decreases towards baseline levels starting from the high values reached during exercise. These trends were qualitatively similar across loading conditions.

**Figure 4:**
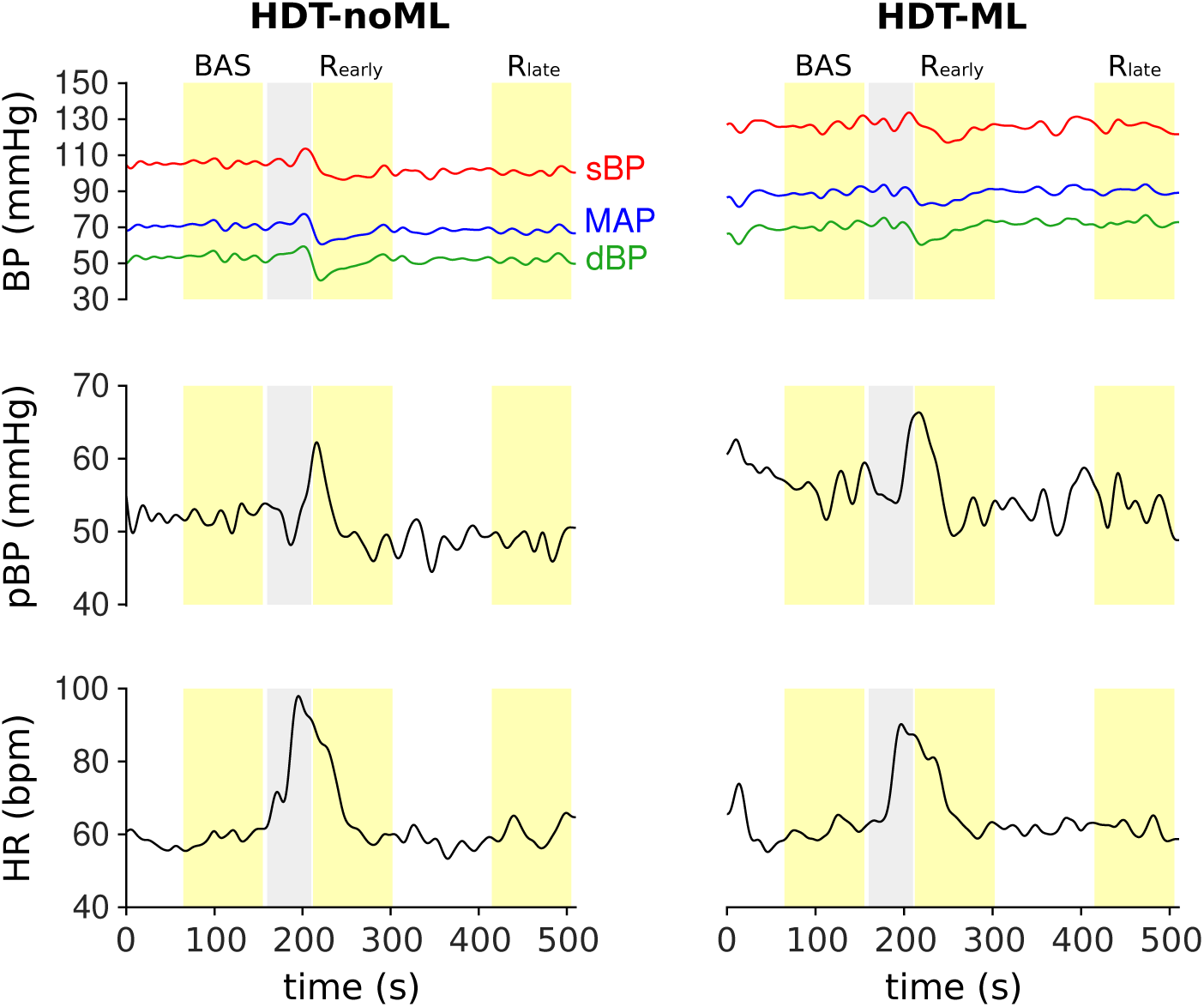
Cardiovascular signals before, during and after dynamic leg-press exercises for one subject. The trends of the signals obtained with antigravity muscle loading (HDT-ML) are qualitatively similar to those obtained without muscle loading (HDT-noML). All the signals exhibit very clear transient dynamics shortly after the exercise bout, and then they stabilize at later time points. Gray band: preparation to exercise (i.e. the subject waits with their legs flexed) and exercise bout. Yellow bands: baseline (BAS), early recovery (R_*early*_, i.e. first 90 seconds after the end of the exercise bout), late recovery (R_*late*_, i.e. 3.5 to 5 minutes after the end of the exercise bout). During baseline and recovery the subject keeps the legs extended. See Sec. 4.2 for details on the protocol.

In order to quantify the transient dynamics observed during early recovery, we defined the following features:

- Minimum values of sBP, dBP and MAP with respect to baseline (ΔsBP_*min*_, ΔdBP_*min*_, ΔMAP_*min*_), to quantify the initial undershoots of these signals;
- Maximum value of pBP with respect to baseline (ΔpBP_*max*_), to quantify its overshoot;
- Maximum value of HR with respect to baseline (ΔHR_*max*_), to quantify the maximum value of this signal at the beginning of the recovery phase.

The comparison between antigravity muscle loading conditions in terms of these features is presented in Fig. 5. The undershoots of sBP and MAP obtained with muscle loading were not significantly different from those observed without muscle loading (ΔsBP_*min*_: +1.1±1.3 mmHg, p=0.429; ΔMAP_*min*_: +2.0±1.3 mmHg, p=0.153). On the other hand, muscle loading resulted in significantly lower values of ΔHR_*max*_ than those obtained without muscle loading (–3.1±1.2 bpm, p=0.024). Finally, results were at the limit of statistical significance for the features ΔdBP_*min*_ (+3.1±1.4 mmHg, p=0.050) and ΔpBP_*max*_ (–3.2±1.5 mmHg, p=0.047). These results suggest that antigravity muscle activity minimally affects the transient dynamics of the cardiovascular signals in recovery from leg-press exercises during HDT.

**Figure 5:**
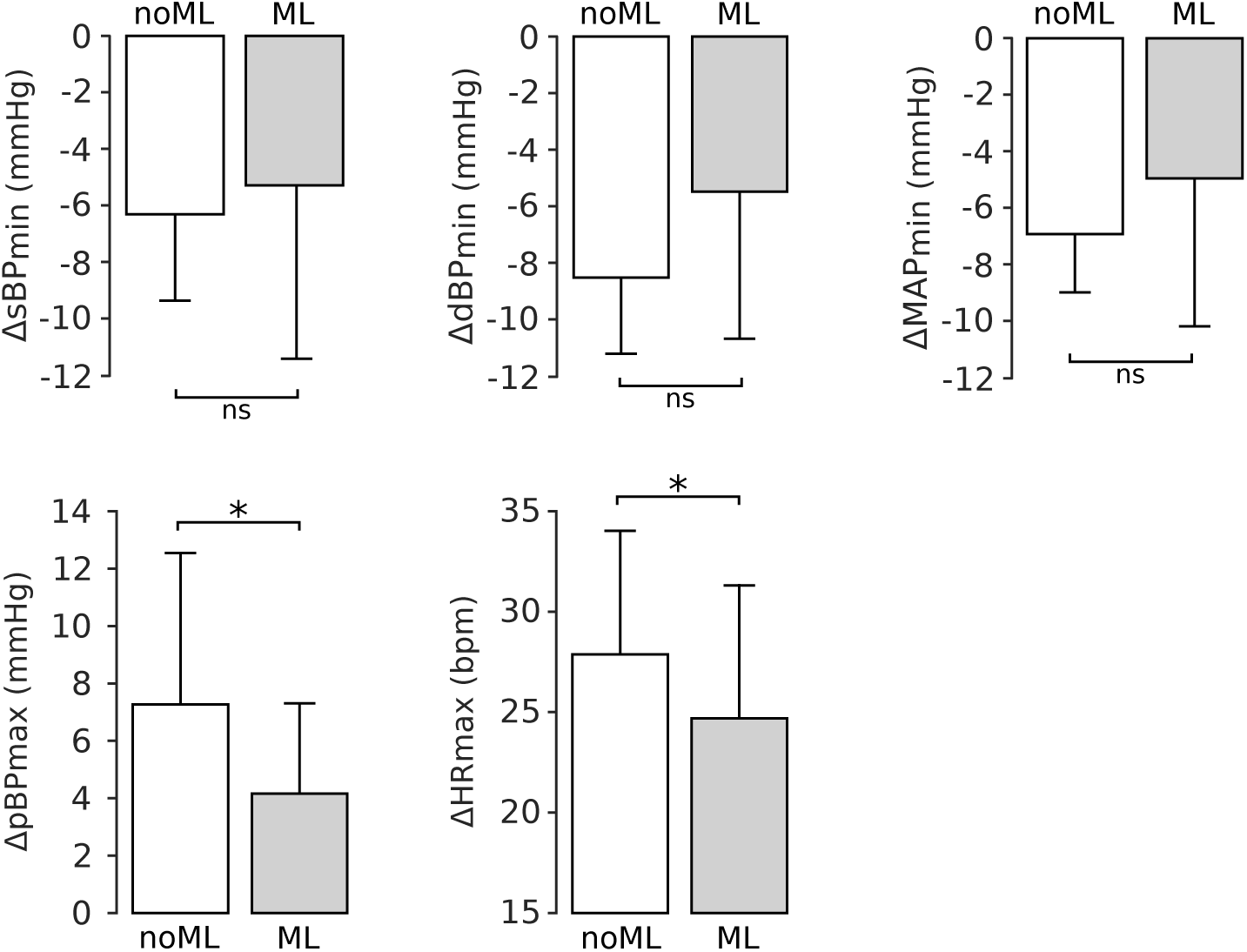
Cardiovascular responses during early recovery with and without antigravity muscle loading for all subjects. Muscle loading resulted in significantly lower pulse BP peaks and heart rate maxima, and did not affect systolic, diastolic and mean arterial pressure. Data are presented as mean±s.d. across subjects. N=17 and 16 subjects contributed to the bars HDT-noML and HDTML respectively (for all signals). *p<0.05.

The values of the cardiovascular signals observed during baseline and late recovery are illustrated in Fig. 6. As already discussed in Sec. 2.2, the values of systolic BP, diastolic BP and MAP obtained with antigravity muscle loading were higher than those obtained without muscle loading. However, there were no significant differences between the values observed during late recovery and those observed at baseline for any signal, both with muscle loading (sBP: +0.8±2.8 mmHg, p=1; dBP: +2.4±2.6 mmHg, p=1; MAP: +1.9±2.4 mmHg, p=1; pBP: –1.6±2.2 mmHg, p=1; HR: –0.3±1.4 bpm, p=1) and without muscle loading (sBP: –0.2±2.7 mmHg, p=1; dBP: +0.3±2.5 mmHg, p=1; MAP: +0.1±2.4 mmHg, p=1; pBP: –0.5±2.2 mmHg, p=1; HR: –0.6±1.3 bpm, p=1). This result is further confirmed by the non-significant interaction term (sBP: p=0.799; dBP: p=0.57; MAP: p=0.623; pBP: p=0.75; HR: p=0.88) between the muscle loading condition (HDT-ML or HDT-noML) and the phase of the experimental session (baseline or R_*late*_) in our statistical model (see Sec. 4). In other words, independently of muscle loading condition, within 5 minutes after the exercise bouts, all the signals returned to values similar to those observed during baseline.

**Figure 6:**
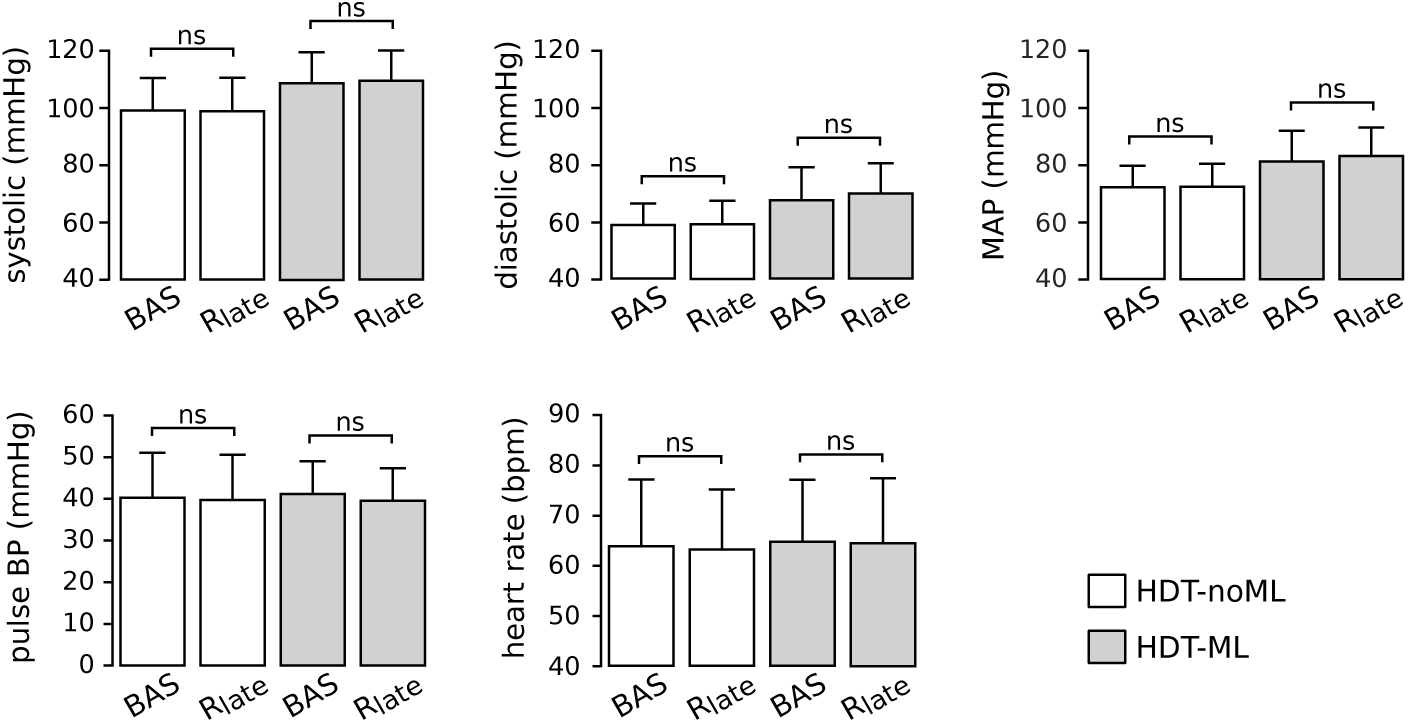
Cardiovascular signals before leg-press exercises and during late recovery with and without antigravity muscle loading. At the end of the recovery phase (R_*late*_), the mean values of all signals returned to values that were similar to those observed before the exercise bouts (baseline, BAS), both with and without antigravity muscle loading (HDT-ML and HDT-noML respectively). Data are presented as mean±± s.d. across subjects. N=17, 17, 16 and 16 subjects contributed to the bars BAS (HDT-noML), R_*late*_ (HDT-noML), BAS (HDT-ML), R_*late*_ (HDT-ML) respectively.

## 3 Discussion

We investigated the influence of antigravity muscles loading on cardiovascular regulation in the absence of orthostatic stress both at rest and after dynamic leg-press exercises. We found that muscle loading caused an increase of BP during HDT at rest, re-establishing orthostatic values of systolic BP. Furthermore, muscle loading affected the transient dynamics of the signals shortly after exercise, with a small reduction of pulse BP and HR peaks relative to baseline, but not the values reached 5 minutes after the end of the exercise bouts. Taken together, these results demonstrate that antigravity muscles activity contributes to cardiovascular regulation in the absence of orthostatic stress.

Two types of neural mechanisms might underlie the influence of muscle activity on cardiovascular regulation: central commands [58, 26], hypothetically originating in motor cortex, and the exercise pressor reflex [21, 45], directly evoked by muscle contraction via activation of mechanoreceptors and metaboreceptors. Both of these mechanisms produce a temporary alteration of the carotid baroreflexes [28], causing the simultaneous increase of BP and of sympathetic discharge and HR. In particular, muscle activity shifts (or “resets”) the carotid-sympathetic baroreflex curve towards higher MAP (rightward shift) and towards higher sympathetic activity (upwards shift). Similarly, it shifts the carotid-cardiac baroreflex curve towards higher MAP (rightward shift) and towards higher HR (upwards shift). As we will discuss below, our results can be explained based on these mechanisms.

Changing posture from upright to HDT (without muscle loading, HDT-noML) caused a reduction of HR, dBP, sBP and MAP. These responses are consistent with previous observations [50, 37, 30], and result from the displacement of fluids from the lower to the upper part of the body. Such a redistribution of body fluids statically stimulates the carotid baroreceptors and increases central blood volume (CBV), thus evoking baroreflex activity that leads to peripheral vasodilatation and a reduction of HR. These factors contribute to a sustained decrease of dBP, sBP and MAP at the heart level [39, 38, 50]. The physiological mechanisms underlying these responses are still under investigation. However, it has been suggested that increasing CBV may reduce the sensitivity of the baroreflexes [5, 49], and may reset their stimulus-response curves towards an operating point characterized by lower MAP and lower HR as well as lower muscle sympathetic nerve activity (MSNA) compared to upright posture [44].

Antigravity muscle loading during HDT (HDT-ML) at rest resulted in significantly higher values of sBP, dBP and MAP, and no change in HR compared to HDT without muscle loading at rest. This finding is consistent with the resetting of the baroreflexes towards higher blood pressure associated with static muscle contraction [41, 28]. However, if the carotid-cardiac baroreflex curve had also relocated upward along the HR response axis as previously observed [28], we would have expected an increase in HR [33]. Since we did not observe such an increase, our results may indicate a pure rightward shift of the carotid-cardiac baroreflex curve preferentially driven by exercise pressor reflex [41]. The lack of change in HR may also result from a low sensitivity of the carotid-cardiac baroreflex due to the elevated CBV in the HDT position [5, 49, 32]. Finally, although we adapted the intensity of muscle loading to the weight of each participant, inter-subject variability may lead to different energy expenditures, which may constitute a potential confounding factor in the analysis. Nevertheless, we are confident of the validity of our results as the minimal alteration of pulse BP and HR after muscle loading is consistent with the expected responses to static exercises, which evoke small changes in cardiac output and a large increase in MAP.

Performing physical exercises elicited well defined after-effects on the cardiovascular signals. The physiological processes that occur in such a recovery period are distinct from those that characterize the resting and the exercising states, and are currently an active topic of research [43]. In the recovery period, the arterial baroreflex curves are reset leftwards to lower operating points in the BP axis and downwards along the response axis, leading to a reduction of HR and sympathetic outflow compared to the exercise period [12, 46, 6]. Accordingly, the elevated HR that we observed at the beginning of recovery slowly returned to baseline values. In addition to this centrally-mediated reduction of sympathetic activity, there are also local mechanisms that cause vasodilation during recovery from exercise [13]. Both these processes may have contributed to the fall of arterial pressure that we observed at the beginning of recovery. This initial reduction of arterial pressure, in concert with the HDT posture that facilitated venous return, may have led to the observed transient increase of pulse BP.

While previous studies demonstrated that prolonged exercise sessions (at least 20 minutes) result in a reduction of arterial blood pressure that may last for hours [14], we found that sBP, dBP and MAP at the end of the recovery period (R_*late*_) were not different from baseline values. This discrepancy may be due to the relatively short duration of the exercise bouts in our study. These short exercise bouts may not elicit the mechanisms leading to sustained vasodilation (e.g. the activation of histamine H1 and H2 receptors [27]), evoking only those that cause immediate postexercise hyperaemia [3].

Antigravity muscle loading influenced post-exercise cardiovascular responses at the beginning of the recovery phase. When subjects had to counteract the force applied by the leg-press device (HDT-ML), we observed slightly lower peak values of pulse BP and HR than in the sessions without muscle loading (HDT-noML) during early recovery R_*early*_ (Fig. 5). The reduction of pulse BP peak is at the limit of statistical significance, and therefore we cannot derive strong conclusions. However, this effect may be due to increased cardiac afterload, which is consistent both with the overall higher values of MAP and with the lower dBP undershoot observed when muscle loading was applied (Fig. 3, 4, 6). The reduction of the maximum value of HR may be due to a milder exercise-induced resetting of the carotid-cardiac baroreflex. It is conceivable that the magnitude of the upwards and rightward displacement of the baroreflex curve associated with dynamic exercise [28] is limited within safe blood pressure ranges. Since baseline MAP with muscle loading was higher than without muscle loading (Fig. 3), then the same exercise intensity may have led to a lower baroreflex displacement to maintain BP within these ranges.

Our results have implications to the cardiovascular deconditioning that occurs during space missions. Similarly to HDT, microgravity causes a headward shift of body fluid that in the short term elicits a reduction of blood pressure and heart rate, and in the long term leads to a reduction of baroreflex sensitivity and ultimately to orthostatic intolerance [30, 20, 16, 53]. In this study, we showed that loading the antigravity muscles during HDT at rest causes an increase of BP that can establish similar values to those associated to orthostatic condition (i.e. earth-like). Similarly, dynamic exercises during HDT can temporarily increase HR and leg blood volume [1]. These findings motivate the development of lower limb wearable devices that can provide astronauts with antigravity muscle loading for extended periods of time, potentially limiting the reduction of blood pressure they experience in the microgravity environment. Long term bed-rest studies and experiments during space missions will be necessary to test whether such a sustained antigravity muscle loading, along with the intense sessions of dynamic exercises that astronauts perform regularly [15, 54], can ameliorate long-term effects of microgravity on the cardiovascular system.

Additional experiments will also be necessary to better understand the physiological bases of the findings reported here. A direct estimation of the baroreflex curves would allow the influence of antigravity muscle loading on baroreflex characteristics during HDT to be quantified. Introducing female subjects could elucidate possible gender-driven modulation of these mechanisms. Furthermore, applying different loading forces could reveal potential intensity-driven modulations. While previous experiments have already considered some of these aspects in orthostatic conditions [40, 47, 4, 9, 2], the results may not directly translate to HDT as this condition is characterized by increased central blood volume and reduced venous pooling when compared to upright posture.

## 4 Methods

### 4.1 Participants

Seventeen male subjects (age: 29.7±3.9 years, weight: 79.2±7.7 kg, height: 1.79±0.06 m, mean±s.d.) volunteered to participate in this study, and signed an informed consent. Subjects had no history of cardiovascular nor musculoskeletal diseases, and were instructed to avoid caffeine and food before the experiment. All procedures were conducted in conformance with the Declaration of Helsinki and were approved by the Ethical Review Committee of Canton Zurich.

### 4.2 Study protocol

The analyses reported in this paper are part of a larger study, which we describe here for completeness and reproducibility. After sensor placement and calibration, we recorded cardiovascular signals during orthostatic stress for 3 minutes (i.e. upright standing posture without movements; ORT condition). Then, we let the subjects lie on the tilted platform (HDT condition) and we secured them to the robotic leg-press device MARCOS (Sec. 4.3). In this posture, they performed nine consecutive experimental sessions, each consisting of three phases: baseline, exercise, and recovery. During baseline, subjects kept their legs extended for 3 minutes. In preparation for the exercise phase, they flexed their legs until reaching the mechanical stop of MARCOS (20 seconds). During the exercise phase, they performed leg-press exercises against the resistance of the device. Finally during recovery, subjects kept their legs extended for 5 minutes.

The intensity of the exercises was defined as a combination of: (1) resistive force of the leg-press, F; (2) frequency of the leg-press movements, f; (3) duration of the exercise phase, T. We tested two values for each of these factors (see here [1] for details), for a total of eight intensities and corresponding experimental sessions. In these sessions, MARCOS did not apply any force during baseline and recovery, so that no muscle contraction was required to maintain leg extension (i.e. head-down tilt without antigravity muscle loading, HDT-noML). In an additional session, the device applied a force of 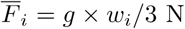 (where *w*_*i*_ is the weight of the *i*-th subject, and *g* = 9.6 *m/s*^2^ is the gravitational acceleration) during all the phases (i.e. head-down tilt with antigravity muscle loading, HDT-ML), and the leg-press exercises were performed at frequency f=0.5 Hz and duration T=30s. The order of these nine sessions was randomized across subjects to avoid biases in the results.

For the purpose of this study, we considered the ORT condition, and only two of the HDT experimental sessions described above: HDT-ML, and HDT-noML with matching exercise intensity, i.e. 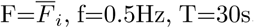. This allowed us to investigate the effects of antigravity muscle loading on the cardiovascular signals during HDT, both at rest (comparing the baseline phases of HDT-ML and HDT-noML, and using ORT as a reference) and during recovery from physical exercise (comparing the relative changes from baseline to recovery observed in HDT-ML and HDT-noML).

### 4.3 MARCOS

MARCOS is a robotic device originally developed to execute leg movements inside a MRI scanner [19]. In this study, we adapted this device to perform subject-driven leg-press exercises on a 6° head-down tilted platform (Fig. 1c). Foot loading is realized by means of pneumatic actuators that can provide a simulated ground reaction force (i.e. opposing resistance to leg extensions) of up to 400 N.

The user is secured to the platforms by means of: (1) shoes that are firmly attached to the foot pedals, (2) a brace that prevents mediolateral movements of the hip, and (3) shoulder straps that prevent sliding in the direction of force. The position of the brace as well as the length of the shoulder straps can be adjusted to the size of the user, allowing us to obtain very similar ranges of motion across subjects (hip: 0° to 40°; knee: 0° to 70°).

### 4.4 Data acquisition and processing

Continuous blood pressure (BP) was measured by means of a CNAP®Monitor 500 (CNSystem Medizintechnik AG, Austria). This system uses the vascular unloading technique to estimate BP information from plethysmographic signals by means of two finger cuffs with integrated infrared light sensors [10]. In order to reduce motion artifacts, we used an adjustable strap to support the arm of the subject in a standardized position, allowing the hand to rest at the level of the heart.

The raw BP signal was low-pass filtered (10Hz, 3rd order Butterworth). Systolic (sBP) and diastolic (dBP) traces were obtained by linear interpolation of the maximum and minimum peaks in the filtered BP wave respectively. Heart rate (HR) was computed as the ratio between 60 s and the time difference between adjacent systolic peaks. The obtained sBP, dBP and HR were further low pass filtered to evaluate their average trends along the experimental sessions (0.05 Hz, 3rd order Butterworth). Pulse blood pressure (pBP) was computed as pBP=sBP-dBP. Mean arterial pressure (MAP) was estimated as MAP = 1/3 sBP + 2/3 dBP.

From these continuous signals, we computed several features to characterize each phase of the experiment. During the orthostatic condition and the baseline phase of the HDT sessions, the signals were stable. Hence, we averaged each continuous trace over the last 90s of recording. On the other hand, during the recovery phase, the signals exhibited an initial transient dynamics and then they stabilized. Therefore, we computed signal-specific features in the first 90 s of recovery (R_*early*_; see Sec. 2.3), and we averaged each signal over the last 90 s of recovery (R_*late*_).

Due to unexpected failures of the acquisition system during the recordings, we had to exclude one subject from the ORT condition and one subject from the HDT-ML condition, resulting in N=16, 17, 16 subjects for ORT, HDT-noML, and HDT-ML respectively.

### 4.5 Statistical analysis

We employed Linear Mixed Effect Models (LMEM) to analyze the cardiovascular signals both at baseline and during the recovery phase, using the nlme package [35] in the R environment. LMEMs allowed us to consider variability at different levels of the dataset (e.g. across conditions and across subjects), and to cope with missing data, thus obtaining maximal statistical power from our dataset. To confirm that our dataset met the assumption of Gaussian distribution and independence of residuals and random effects [36], we visually inspected the distributions using qq-plots and histograms, and we employed Shapiro-Wilk Normality tests. After fitting the LMEMs, we tested our specific hypotheses of interest by performing post-hoc tests and using Bonferroni corrections to adjust the obtained p-values. Note that because of this correction, the adjusted p-values can be equal to 1. We considered tests to be statistically significant if their p-values were lower than the 0.05 significance level.

To assess the effect of antigravity muscle loading at rest and at the end of the recovery phase (R_*late*_), we fit a LMEM for each signal, using the average value of the signal across the 90 s window of interest (see Sec. 4.4) as the dependent variable, and the following independent variables: subject weight (continuous variable), the phase of the experimental session (a factor with levels *baseline* and *recovery*), and the muscle loading condition (a factor with levels *ML* and *noML*). Finally, we considered the interaction term between phase and loading to analyze the potentially different effects of muscle loading at baseline (rest condition) and during R_*late*_. In order to take inter-subjects variability into account, we considered subject identity as a random effect on the model intercept, obtaining a more powerful version of a repeated measure ANOVA. We then performed post-hoc tests to evaluate whether: (1) the baseline values of the signals with muscle loading were different from those without muscle loading; (2) the R_*late*_ values of the signals were different from baseline, both with and without muscle loading.

To evaluate the differences between the signals observed during orthostatic stress (ORT) and those recorded during head-down tilt at rest with and without muscle loading (HDT-ML and HDT-noML respectively), we fit a LMEM for each signal using the average value of the signal as the dependent variable, and the experimental condition as the independent variable. We also considered subject identity as a random effect on the model intercept. We then performed post-hoc tests to assess if the values of the signals obtained during each HDT condition were different from those obtained during ORT.

Finally, to evaluate the effect of muscle loading on the transient dynamics of the signals during early recovery (R_*early*_), we fit a LMEM for each signal using the signal-specific feature (see Sec. 2.3) as the dependent variable, and subject weight and muscle loading condition as the independent variables. We also considered subject identity as a random effect on the model intercept. We performed post-hoc tests to evaluate if the values of the features obtained with muscle loading were different from those obtained without muscle loading. We report the results of each post-hoc test as mean±s.e., and the corresponding p-values.

## Data availability

The dataset and the code used for the current study are available in the OSF repository, https://osf.io/fgvh3/.

## 5 Acknowledgements

This research was funded by the STAMAS Project of the EU Seventh Framework Programme FP7/2007-2013 under Grant 312815 STAMAS. The authors would like to thank Dr. Klamroth-Marganska for the medical advices, Mrs. Maya Kamber for the support to obtain ethical approval, and Prof. Matthew Tresch for providing feedback on the manuscript. Finally, we thank Mr. Michael Herold-Nadig, Dr. Laura Marchal-Crespo and Dr. Juan Pablo Carbajal to help setting up and testing the robot MARCOS.

## 6 Author contribution

CA and RR designed the experiments. CA conducted the experiments, analyzed the data, and wrote this manuscript. AST contributed to the data analysis and the algorithms, and provided technical support with the equipment. CA, AST and RR revised the manuscript.

